# The mechanism of bacterial defense system DdmDE from *Lactobacillus casei*

**DOI:** 10.1101/2024.10.09.617351

**Authors:** Pingping Huang, Purui Yan, Lijie Guo, Wenying Fei, Zhaoxing Li, Jingxian Liu, Jianping Kong, Yue Yao, Meiling Lu, Yibei Xiao, Meirong Chen

## Abstract

Bacteria has developed a diverse array of defense mechanisms to protect against invasion by mobile genetic elements. Recent study identified a bacterial defense module DdmDE system which encodes a helicase-nuclease fusion protein DdmD and a putative prokaryotic Argonaute-like protein DdmE, imposing fitness advantages to the bacteria by eliminating invasive plasmids. However, the mechanistic basis of how DdmDE system detects and degrades plasmids is not fully understood. Here, by studying the DdmDE system from *Lactobacillus casei* (LcDdmDE), we found that LcDdmD is able to degrade ssDNA and nick plasmids in the presence of Mn^2+^, and it exhibits 5’-3’ DNA helicase activity in a ssDNA length-dependent mechanism. Meanwhile, LcDdmD serves as a sensor that utilizes DNA guide to recognize target DNA. We determined the cryo-EM structures of LcDdmD dimer bound with fork DNA, guide/target DNA-bound LcDdmE, and the complex of LcDdmDE-bubble DNA in intermediate state as well as active state. Together with functional analysis, we revealed the working mechanism of LcDdmDE system. In such a scenario, guided by ssDNA, LcDdmE recruits auto-inhibited LcDdmD dimer loading onto DNA target. Through substantial conformational changes, LcDdmD dimer dissociates into active monomer and unwind duplex for plasmid degradation. Our study provides structural insights into the mechanism of DdmDE, presenting pAgo-directed plasmid degradation by the allosterically regulated helicase-nuclease.

---

Prokaryotes have evolved a multitude of defense systems that confer immunity against invasive mobile genetic elements ^[1–5]^. A two-gene DNA defense module DdmDE system is found in Vibrionaceae and Lactobacillaceae family and imposes fitness advantages to the bacteria by eliminating invasive plasmids ^[6]^. DdmD encodes a fusion protein comprising an N-terminal superfamily 2 helicase and a C-terminal PD-(D/E)XK superfamily nuclease, while DdmE is a putative prokaryotic Argonaute-like protein that frequently engages in bacterial immunity. Although recent studies described the function and structure of DdmDE from *Vibrio cholerae* ^[6–9]^, the mechanistic basis by which the DdmDE system detects and degrades plasmids is not fully understood. In particular, the process by which DdmD dimer dissociates and how the DNA is captured by the nuclease domain for degradation remain elusive.

Here, we focus on DdmDE system from *Lactobacillus casei* (LcDdmDE), which shares only 14% sequence identity with the reported *Vibrio cholerae* DdmDE (VcDdmDE) ^[6–9]^. LcDdmD is able to degrade ssDNA and nick plasmids in the presence of Mn^2+^ (Fig. 1a; Supplementary information, Fig. S1a), and it exhibits ATP-dependent 5’-3’ DNA helicase activity (Fig. 1a). Unlike VcDdmE which only recognizes 5’-phosphorylated guide DNA (5’-P gDNA), LcDdmE uses both 5’-P and 5’-OH gDNA to recognize target DNA (Fig. 1b), with 5’-P gDNA more preferred. DdmE also exhibits a slight binding affinity for guide DNA-target RNA and guide RNA-target RNA, albeit at significantly lower levels (Supplementary information, Fig. S1b). Moreover, compared with VcDdmE that prefers gDNA shorter than 14 nucleotides ^[7,8]^, LcDdmE demonstrated comparable binding affinity across various lengths of gDNA spanning from 14 to 21 nucleotides (Supplementary information, Fig. S1c). Aligning with the absence of a canonical DEDX catalytic tetrad in the PIWI domain, LcDdmE lacks the ability to catalyze DNA-guided target DNA cleavage (Supplementary information, Fig. S1d). These observations collectively suggest that DdmE likely serves as a sensor to recognize invasive elements via a DNA-guided DNA targeting mechanism, while DdmD probably functions as the effector responsible for eliminating the target DNA.

**Fig. 1.**
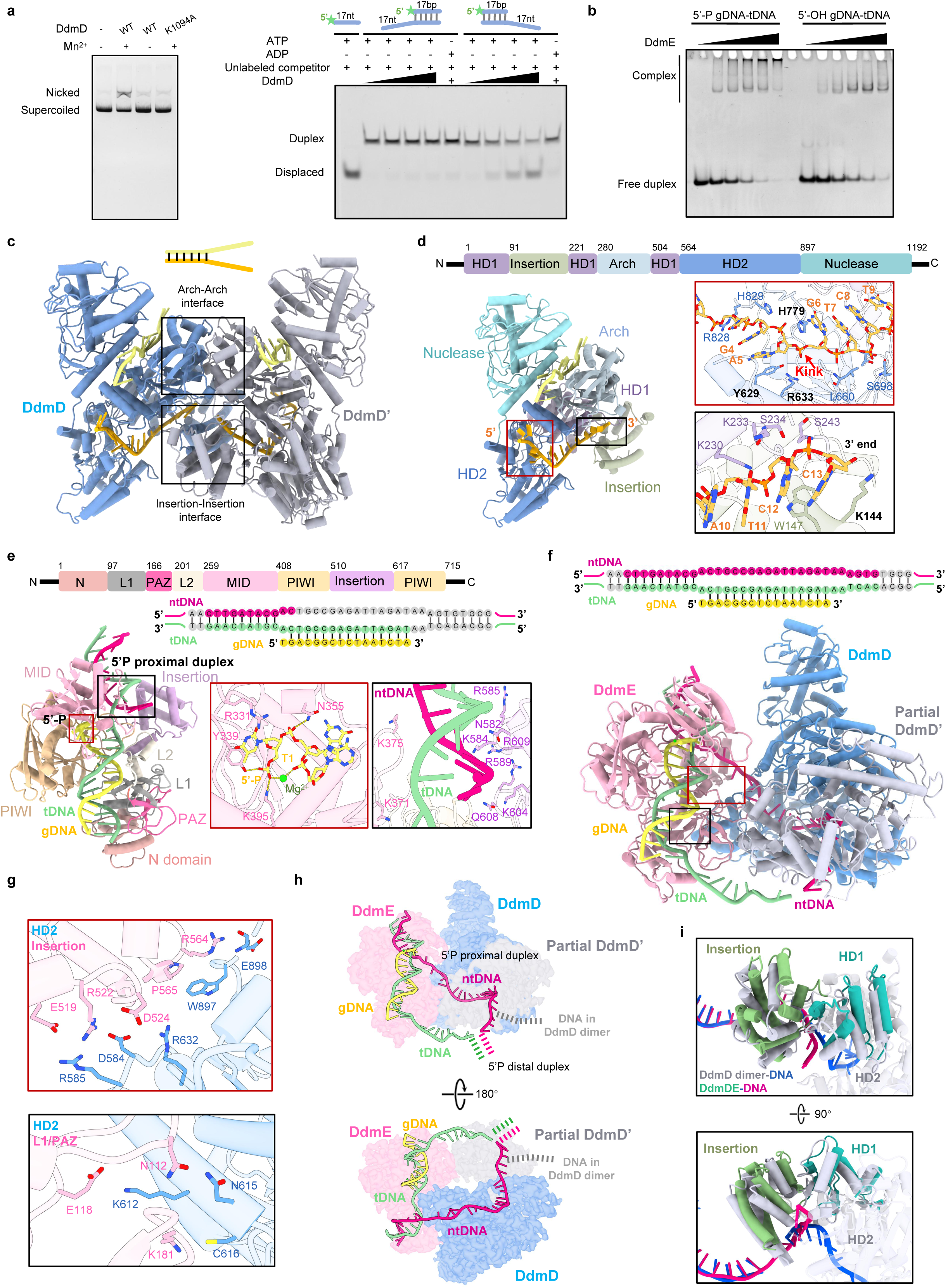
Functional analysis and cryo-EM structures of DdmDE system from *Lactobacillus casei*. a. The characterization of nuclease (left) and helicase (right) activity of LcDdmD. b. The guide and target preference of LcDdmE. c. The cryo-EM structure of LcDdmD in complex with fork DNA. d.The recognition of DNA by DdmD. The second protomer of LcDdmD dimer with the bound DNA are omitted for clarity. The domains of LcDdmD are colored according to the primary structure at top. The two insets highlight the kinked nucleotides (top) and the 3’ end of ssDNA (below), respectively. e. The cryo-EM structure of LcDdmE in complex with guide and bubble DNA. The domains of DdmE are colored according to the primary structure at top; The DNA sequences used in structure are shown, with guide DNA colored in yellow, target strand of bubble DNA in green, non-target strand in magenta, invisible nucleotides in grey. The middle inset shows the recognition of target strand of bubble DNA via guide-target complementarity; The right inset display the interactions between the insertion domain of DdmE and the 5’P-proximal duplex. f. The cryo-EM structure of LcDdmDE-gDNA-bubble DNA. The fully visible promotor of DdmD is colored in blue, partial promotor in grey, and DdmE is colored in pink. The interface between DdmD and DdmE are indicated with square. g. The interactions between LcDdmD and LcDdmE. h. The representation of DNA bound by LcDdmDE. The route of bubble DNA in DdmD-DdmE complex highlights the delivery of non-target DNA to DdmD by DdmE via guide-mediated target binding. i. The conformational changes of the partial protomer of DdmD prior to activation and the opening of the exit for the 3’ end of the non-target DNA.

To unravel the mechanism of DNA unwinding by LcDdmD, we determined the cryo-EM structure of fork DNA-bound LcDdmD at 3.0 Å resolution (Supplementary information, Figs. S2, S3). LcDdmD forms a dimer (Fig. 1c), with each protomer consisting of an N-terminal HD1(1-90, 221-279, and 504-563 aa) separated by an Insertion (91-220) and an Arch domain (280-503 aa), a HD2 domain (564-896 aa) and a C-terminal nuclease domain (897-1192 aa) (Fig. 1d). These elements assemble at Insertion and Arch domain to form a homodimer with a 2-fold symmetry (Fig. 1c). The Insertion and Arch domain are also responsible for dimerization of VcDdmD, however their fold and the dimer interface are not very conserved between LcDdmD and VcDdmD (Supplementary information, Fig. S4). Within the highly positively charged DNA binding channel composed by HD1 and HD2, we modeled 13 nucleotides of the 15-nt long 3’-overhang of the fork DNA in each protomer. The nucleotides stack one after the other (Fig. 1d), except for a kink observed at the fourth and fifth nucleotide positions due to stacking and electrostatic interactions with Y629, H779, and R633 (Fig. 1d). Of note, the DNA 3’ end halts at the dimer interface, with K144 stacking with the final nucleotide (Fig. 1d). This observation, together with the structural similarity of the helicase regions between LcDdmD and DinG, a monomeric 5’-3’ direction DNA helicase ^[10,11]^, suggests that the dimer interface of DdmD potentially obstructs DNA translocation, and dissociation into monomer may be imperative for DdmD activation. Intriguingly, we also observed a class of DdmD monomers in complex with fork DNA, likely representing an active state, albeit with limited resolution due to preferred orientation (Supplementary information, Fig. S5).

We speculated that the extension of ssDNA towards the dimer interface might trigger dimer dissociation to activate DdmD. In agreement, the electrophoresis mobility shift assay showed an increased amount of DdmD monomer with the extension of ssDNA overhang range from 10 to 20 nt, while increase to 25 nt would not further enhance dimer dissociation (Supplementary information, Fig. S6a). As a result, the extension of ssDNA facilitates DNA unwinding by DdmD (Supplementary information, Fig. S6b). These together indicate that LcDdmD is activated through dimer dissociation triggered by ssDNA of an appropriate length. This length-dependent activation by ssDNA was also observed in VcDdmD ^[8, 9]^. Interestingly, we observed a continuous density of ssDNA within a positively charged groove sandwiched by the nuclease domain and the HD2 domain (Supplementary information, Fig. S7a). This ssDNA likely originates from the 5’-overhang of the fork DNA, mimicking the ssDNA protruding from helicase domains during duplex unwinding by DdmD. We also observed the density of ADP at the interface of HD1-HD2, embraced by residues T31, K33, S34, Y35, and R853 (Supplementary information, Fig. S7b), resembling the ADP recognition in DinG ^[10]^.

To elucidate the mechanism of how LcDdmE recognizes the target DNA and recruits LcDdmD for DNA degradation, we reconstituted LcDdmDE complex using a 16-nt 5’-P gDNA and a bubble dsDNA target. The complex was subjected to cryo-EM analysis, and after iterative rounds of 3D classification and refinement, we successfully determined the structures of LcDdmE-gDNA-bubble DNA and LcDdmDE-gDNA-bubble DNA at a resolution of 3.6 Å and 3.3 Å, respectively (Supplementary information, Figs. S2, S3).

LcDdmE resembles typical pAgo structure ^[12]^, consisting of an N-terminal domain (1-96 aa), L1(97-165 aa), a short PAZ domain (166-200 aa), L2 (201-258 aa), MID domain (259-407 aa), and PIWI domain (408-509 and 617-715 aa). Notably, a unique insertion domain (510-616 aa) without homologous structure protrudes from the PIWI domain (Fig. 1e). Fourteen base pairs of the guide-target duplex spanning nucleotide 2-15 are embedded within the highly positively charged channel formed by characteristic two-lobe architecture of Argonaute, with first ten nucleotides exhibiting extensive electrostatic interactions with the residues in channel (Supplementary information, Fig. S7c). Similar to other prokaryotic Ago-guide structures ^[12]^, the first nucleotide of the gDNA flips and anchors in the MID pocket. The R331 and N355 interact with the base moiety of the first nucleotide and the phosphate of the second nucleotide, respectively, with the 5’-P coordinating with magnesium and interacting with Y339 and K395 (Fig. 1e). The substitution for 5’-OH of guide DNA is likely to weaken the interaction considering the longer distance and the weaker electrostatic ability between Y339/K395 and 5’-OH, which may result in slight weaker binding observed in EMSA. The 5’-P proximal duplex region of bubble DNA is hold by insertion domain and MID domain through electrostatic interactions with a cluster of positive charged residues, including K371, K375, N582, K584, R585, R589, K604, Q608, and R609 (Fig. 1e). However, the non-target loop of bubble and the distal duplex are not visible due to flexibility. These findings suggest that DdmE may sense and stabilize the end of the duplex and facilitate loading of DdmD onto the adjacent bubble loop.

The LcDdmDE-gDNA-bubble target complex structure reveals an asymmetric stoichiometry of 2:1 DdmD:DdmE (Fig. 1f). However, only one protomer of LcDdmD could be fully assigned in this complex. In contrast, the other protomer exhibits weak density, with only the HD1, Insertion, and Arch domains, which contribute to the dimer interface, being modeled (Fig. 1f). These observations suggest that this structure may represent an intermediate state prior to activation, in which LcDdmD dimer is undergoing dissociation. The structure reveals direct interactions between LcDdmD and LcDdmE, primarily involving the HD2 domain from one LcDdmD protomer and the insertion/L1/PAZ domain of DdmE, resulting in a buried area of 1073 Å^2^. At the HD2^DdmD^-Insertion^DdmE^ interface, D584, R585, R632, and E898 from DdmD form hydrogen bonds with R522, E519, D524, and R564 from DdmE, respectively (Fig. 1g), and W897^DdmD^ stacks with P565^DdmE^; On the other side, the HD2-L1/PAZ contacts involving K612^DdmD^, N615^DdmD^, C616^DdmD^, N112^DdmE^, E118^DdmE^, and K181^DdmE^ further reinforces the DdmDE complex (Fig. 1g). The remaining Insertion domain of the partial DdmD protomer also shows weak contact with the L1 domain, which may be transient during DNA translocation (Supplementary information, Fig. S8). Although both LcDdmDE and VcDdmDE utilize Insertion^DdmE^ domain and HD2^DdmD^ domain for their interactions, there are notable differences in the fold of insertion domain and its binding interface with HD2 between the two complexes ^[7–9]^ (Supplementary information, Figs. S4b, S4c, S9). These distinctions suggest that DdmE, by employing a unique Insertion domain, plays a role in recruiting other than activating DdmD.

Through the aforementioned interactions, DdmE recruits DdmD onto the non-target DNA strand of bubble DNA. The non-target loop, extending from 5’ P proximal duplex bound by DdmE, traverses into the first DdmD protomer in a direction akin to that of DdmD-fork DNA (Fig. 1h). However, exiting from the first protomer, albeit with weak density, the non-target DNA changes direction and extends along HD1-Insertion interface of the partial dissociated DdmD protomer (Figs. 1h, i). The protruding non-target DNA eventually merges with the target DNA to form a duplex (the density of duplex is traceable but not amenable for modeling) (Fig. 1h). This structure captures the handover of the non-target loop of the bubble DNA to DdmD by DdmE.

Besides the release of Nuclease and HD2 domains in the second DdmD protomer, superimposition of LcDdmDE-gDNA-bubble DNA and LcDdmD dimer-fork DNA also reveals substantial shift in the insertion and HD1 domains of the partial DdmD protomer, resulting in the opening of the exit for the 3’ end of the non-target DNA (Fig. 1i). The movements of Insertion domain weaken the DdmD-DdmD interface, reducing the buried area from 2200 Å to 1935 Å, which may eventually lead to disruption upon the initiation of DNA translocation. Indeed, through deep 3D classification, a fully dissociated DdmD with a 1:1 DdmD:DdmE stoichiometry was resolved at a low resolution (Supplementary information, Fig. S10). In this structure, following the release of the HD1 and insertion domains, the protruding non-target and target DNA are no longer visible, exhibiting a structure similar to those reported for VcDdmDE ^[7,8]^. In addition, the two DdmDE structures also exhibit notable differences in the nuclease domain of DdmD, which shifts 10 Å away from DdmE upon complete dissociation of the dimer. This displacement may indicate a reorientation of the nuclease domain for plasmid degradation during activation; alternatively, it could represent random snapshots of the flexible nuclease domain in various states, as the nuclease domain consistently displays relatively weak and diffuse density in each of our structures.

In summary, our study, in line with parallel investigations on VcDdmDE ^[7–9]^, provides valuable insights into the mechanism of the DdmDE defense system: DdmE senses the invasive plasmid directed by guide DNA, which may be generated by RecBCD and DdmD. Subsequently, DdmE recruits DdmD dimer onto the bubble region of plasmid via DNA-guided DNA targeting, which activates DdmD for plasmid degradation through step-by-step dissociation of DdmD dimer in a ssDNA-dependent manner (Supplementary information, Fig. S11). The mechanism of DdmDE reminiscent the type I CRISPR-Cas3 system, in which the effect complex cascade recruits helicase-nuclease fusion Cas3 upon target DNA recognition ^[13,14]^. Further investigations are necessary to unravel the dynamic process of DNA translocation and degradation, cooperatively mediated by the helicase and nuclease domains of DdmD in the presence of DdmE.

## ACKNOWLEDGEMENTS

We thank the Instrument Analysis Center (IAC) at Shanghai Jiao Tong University for cryo-EM data collection. This work is supported by National Key Research and Development Program of China (2023YFC3402300), STI2030-Major Projects (2021ZD0203400), National Natural Science Foundation of China (32271330; 82321005; 82304614).

## AUTHOR CONTRIBUTIONS

M.C. and Y.X. directed the research. P.H., P.Y., L.G., W.F., J.L., J.K. and Y.Y. performed experiments. M.C., P.H. and Z.L. determined cryo-EM structures. M.C., Y.X., P.H., P.Y., L.G. and M.L. analyzed data. M.C. and Y.X. wrote the manuscript.

## ADDITIONAL INFORMATION

Supplementary information is available online.

## DATA AVAILABILITY

The atomic coordinates of LcDdmD-fork DNA, LcDdmE-gDNA-bubbleDNA, and LcDdmDE-guide-bubbleDNA have been deposited in the RCSB PDB under accession number 9IX4, 9IW3 and 9IXM, respectively, and the corresponding cryo-EM maps have also been deposited in the EMDB under accession number EMD-60964, EMD-60944, and EMD-60973.

---

**Supplementary information, Fig. S1.**
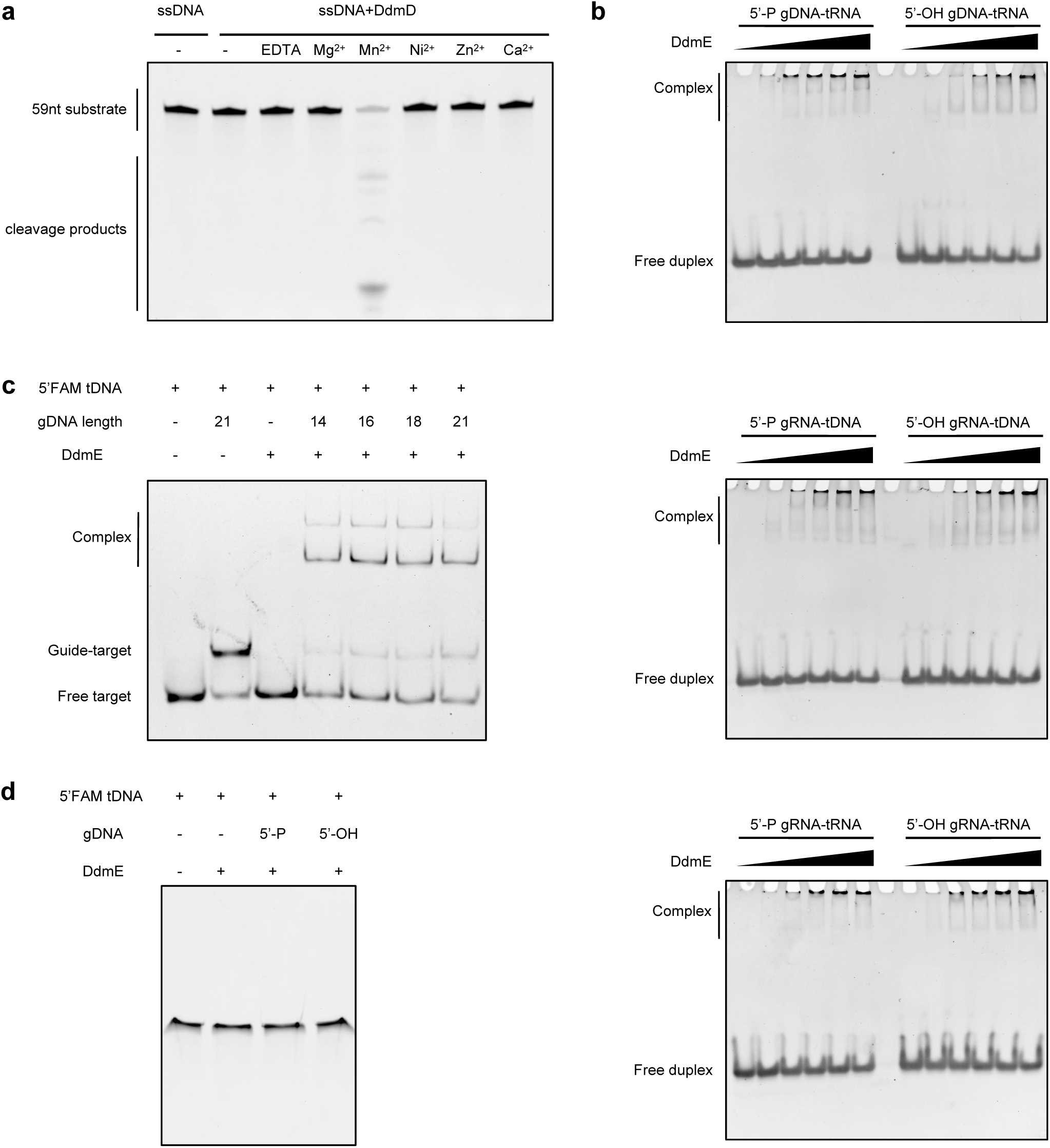
The biochemical characterization of LcDdmDE. a. The manganese-dependent cleavage of ssDNA catalyzed by LcDdmD. b. The EMSA analysis of DdmE binding with various guide-target. c. The EMSA analysis shows the comparrable binding of LcDdmD to DNA guide with various lengths spanning form 14 to 21nt. d. The analysis of target cleavage by LcDdmE. gDNA, guide DNA; gRNA, guide RNA; tDNA, target DNA; tRNA, target RNA.

**Supplementary information, Fig. S2.**
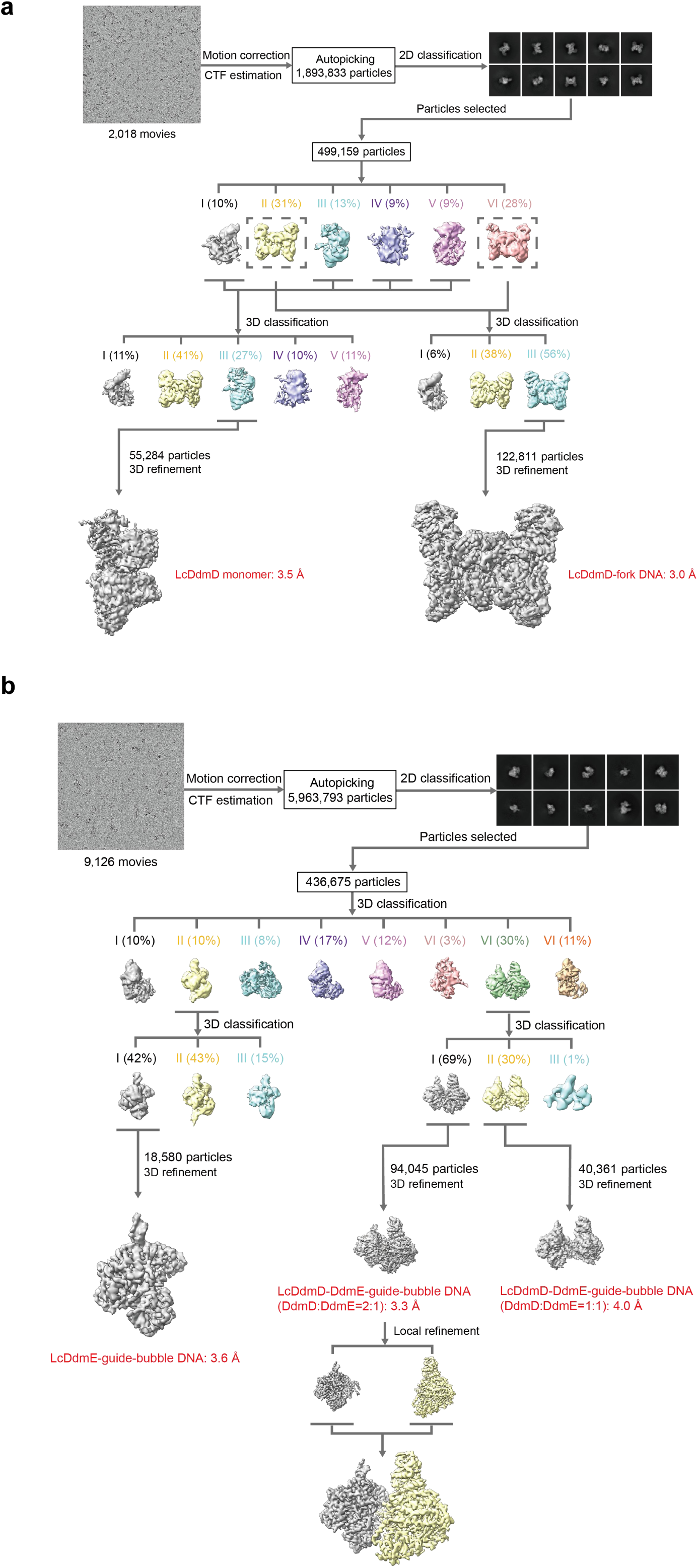
The workflow of cryo-EM data processing.

**Supplementary information, Fig. S3.**
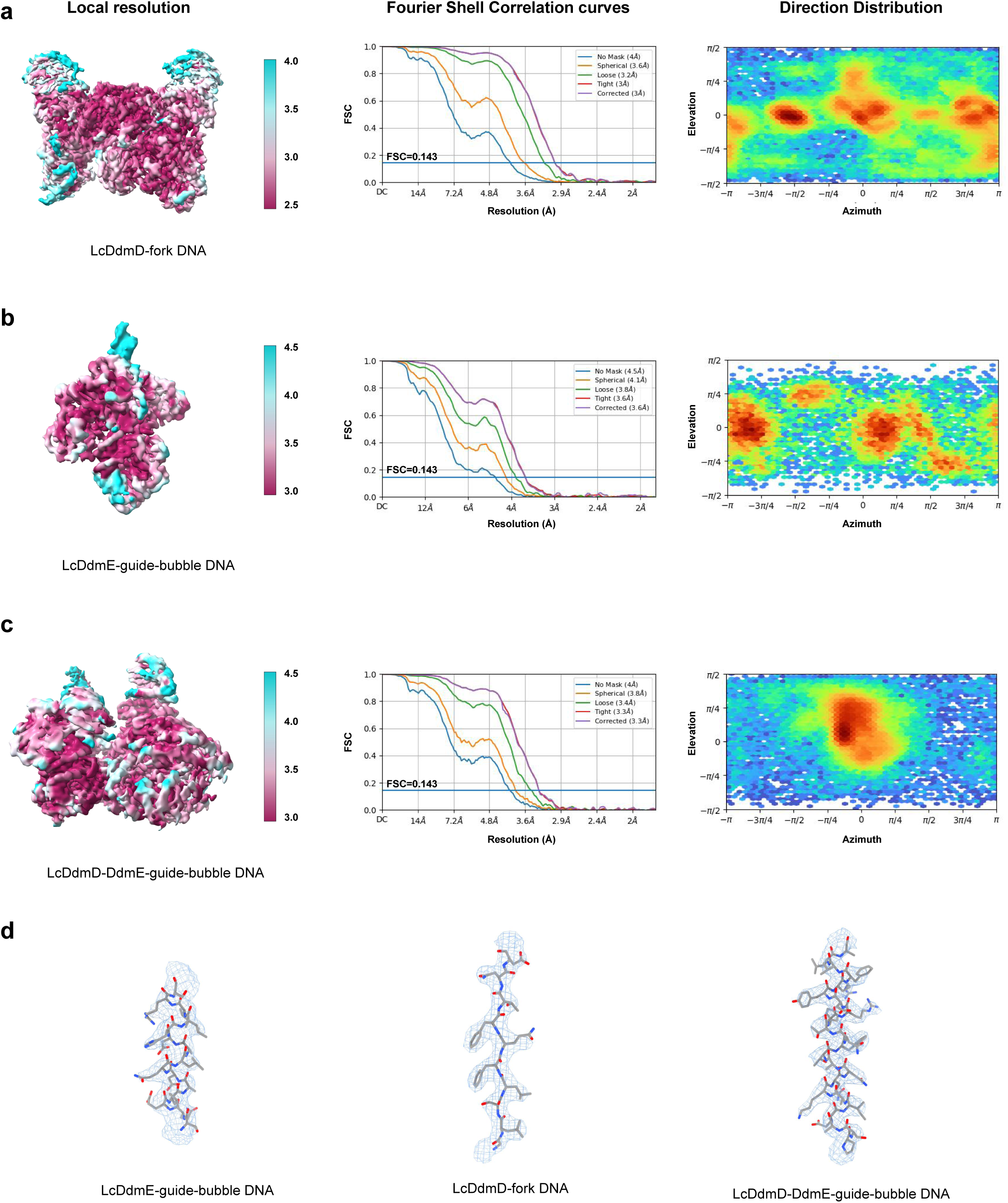
The representation of Cryo-EM map by local resolution. The Cryo-EM maps colored by local resolutionand the Gold-standard Fourier shell correlation curves for LcDdmD-fork DNA (a), LcDdmE- guide-bubble DNA (b), and LcDdmD-DdmE-guide-bubble DNA (c). The cryo-EM density maps with fitted models (d).

**Supplementary information, Fig. S4.**
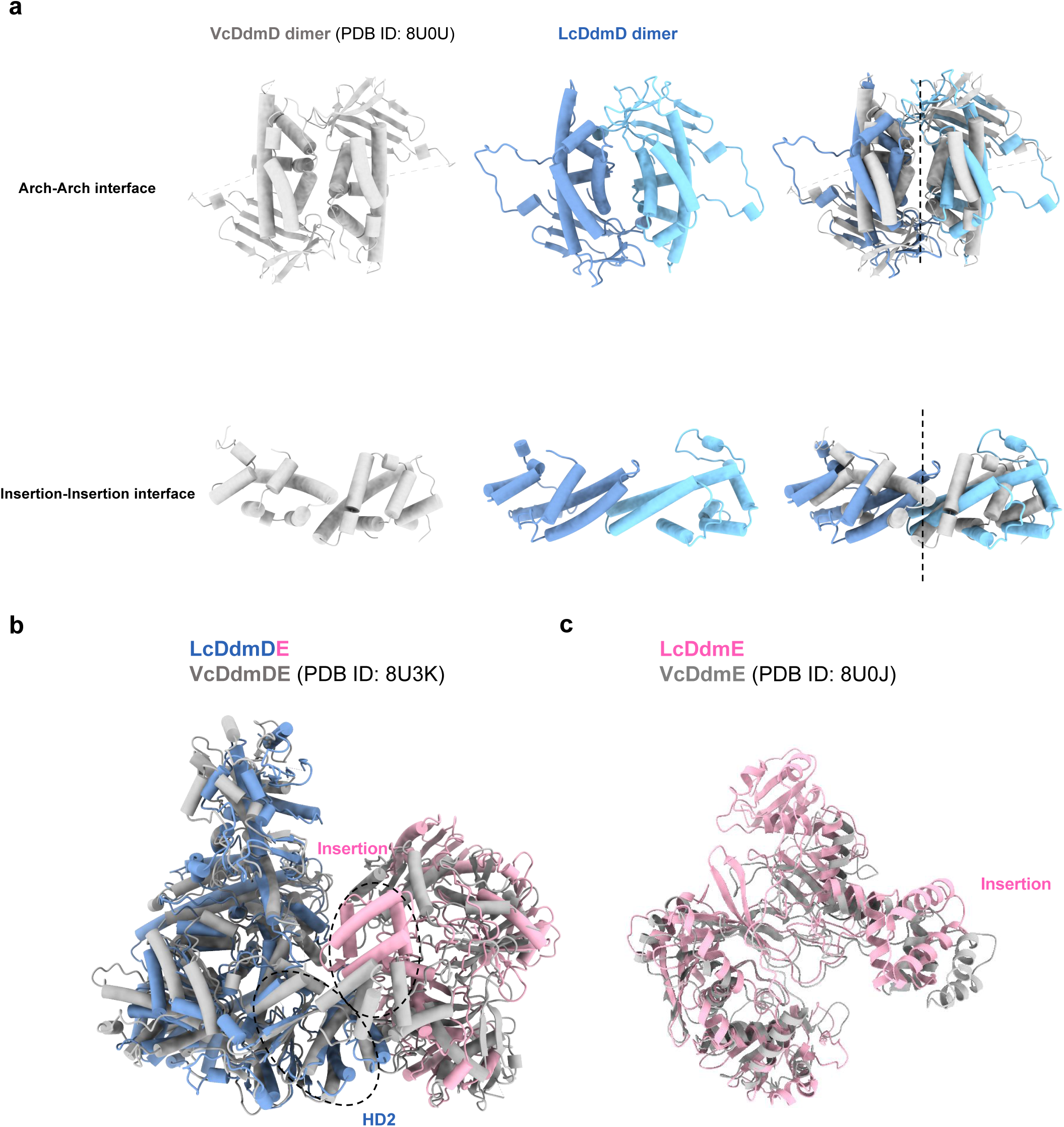
Structural comparison of LcDdmD and VcDdmD. a. Superimposition of LcDdmD and VcDdmD shows the divergent fold at dimer interfaces. b.Superimposition of LcDdmDE onto VcDdmDE shows distinct DE interface involving HD2 and Insertion domains. c. Superimposition of LcDdmE onto VcDdmE reveals similar fold except for the Insertion (N domain is invisible in VcDdmE).

**Supplementary information, Fig. S5.**
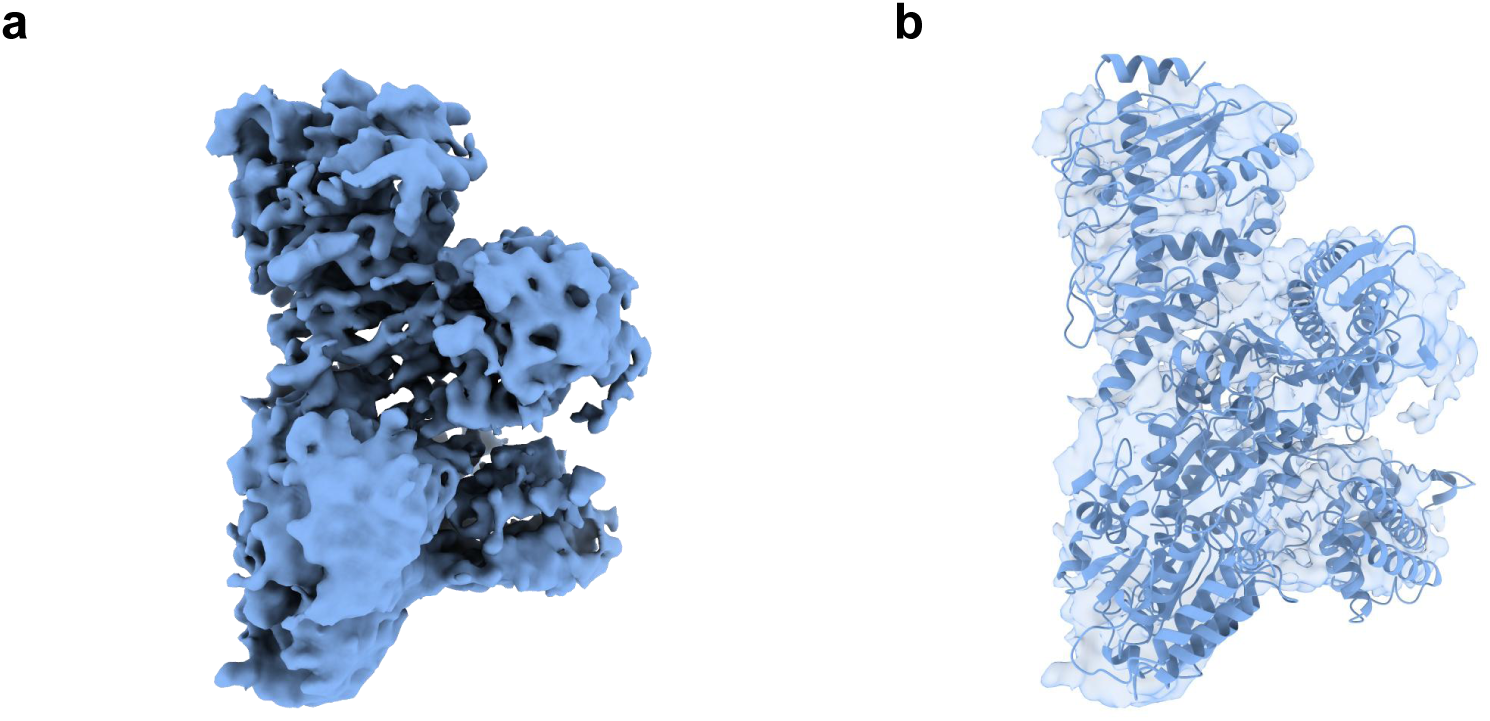
The monomeric LcDdmD isolated from the sample of DdmD-fork. a. The Cryo-EM map of monomeric DdmD shows preferred orientation. b. The model of monomeric DdmD fit into map.

**Supplementary information, Fig. S6.**
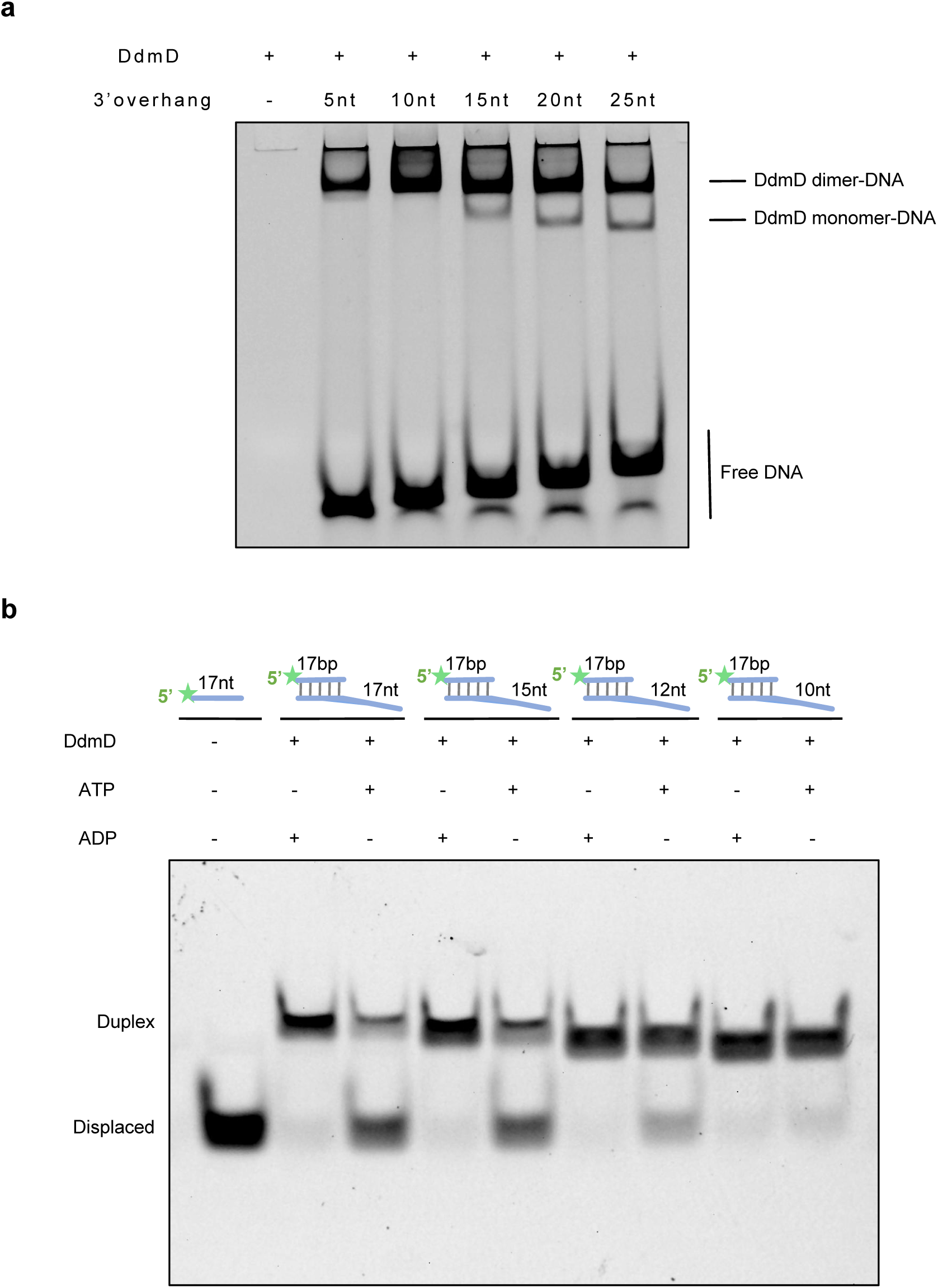
The length-dependent activation of LcDdmD by ssDNA. a. The ssDNA length-dependent LcDdmD monomer formation. b. The ssDNA length-dependent activation of LcDdmD helicase.

**Supplementary information, Fig. S7.**
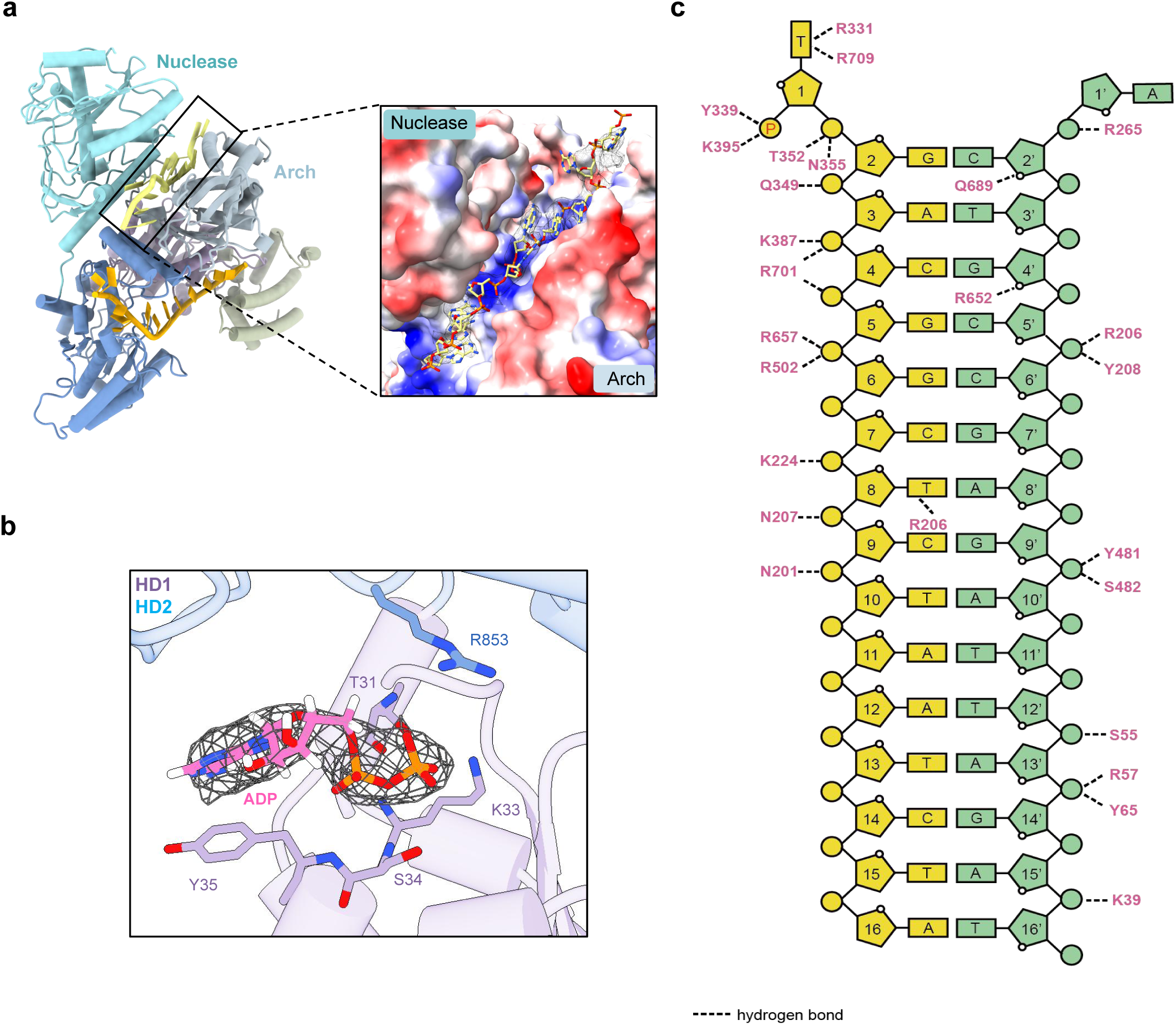
The Interaction between LcDdmD/DdmE and nucleic acids. a. The ssDNA observed in the positive charged channel between Nuclease and Arch domain. b. ADP bound at interface of HD1 and HD2. c. The schematic diagram of interactions between LcDdmE and guide DNA-target DNA.

**Supplementary information, Fig. S8.**
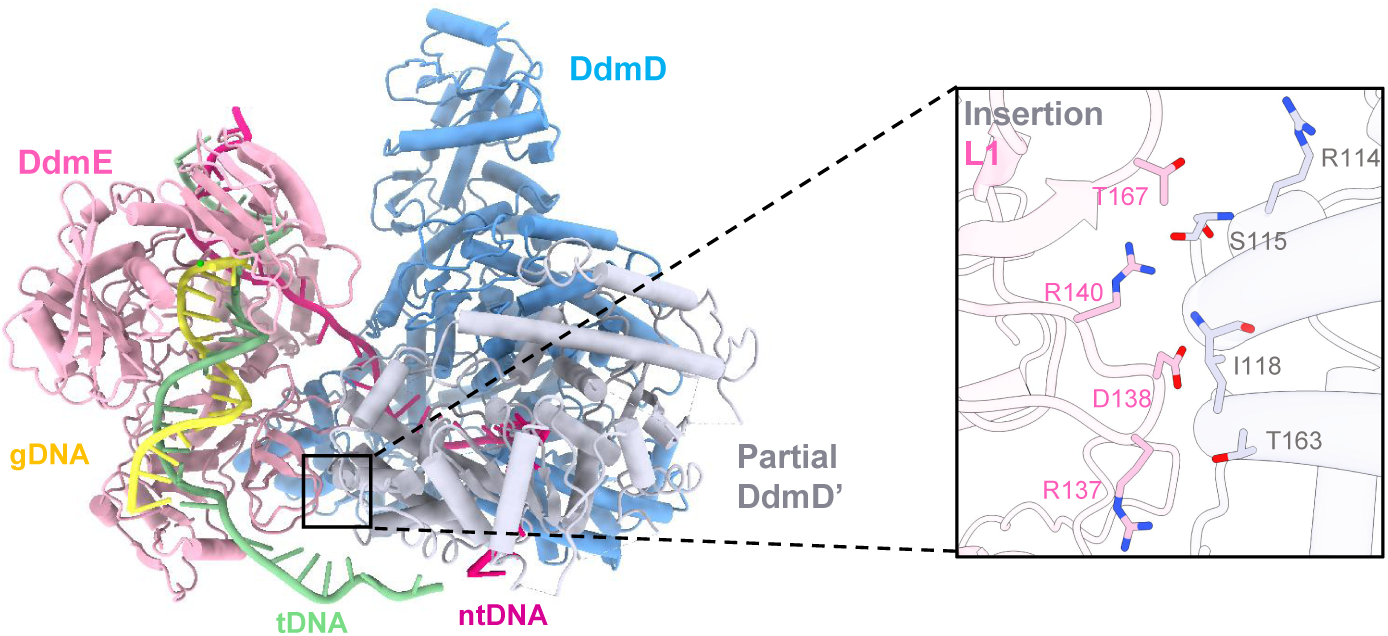
The interaction between LcDdmE and the partial protomer of LcDdmD.

**Supplementary information, Fig. S9.**
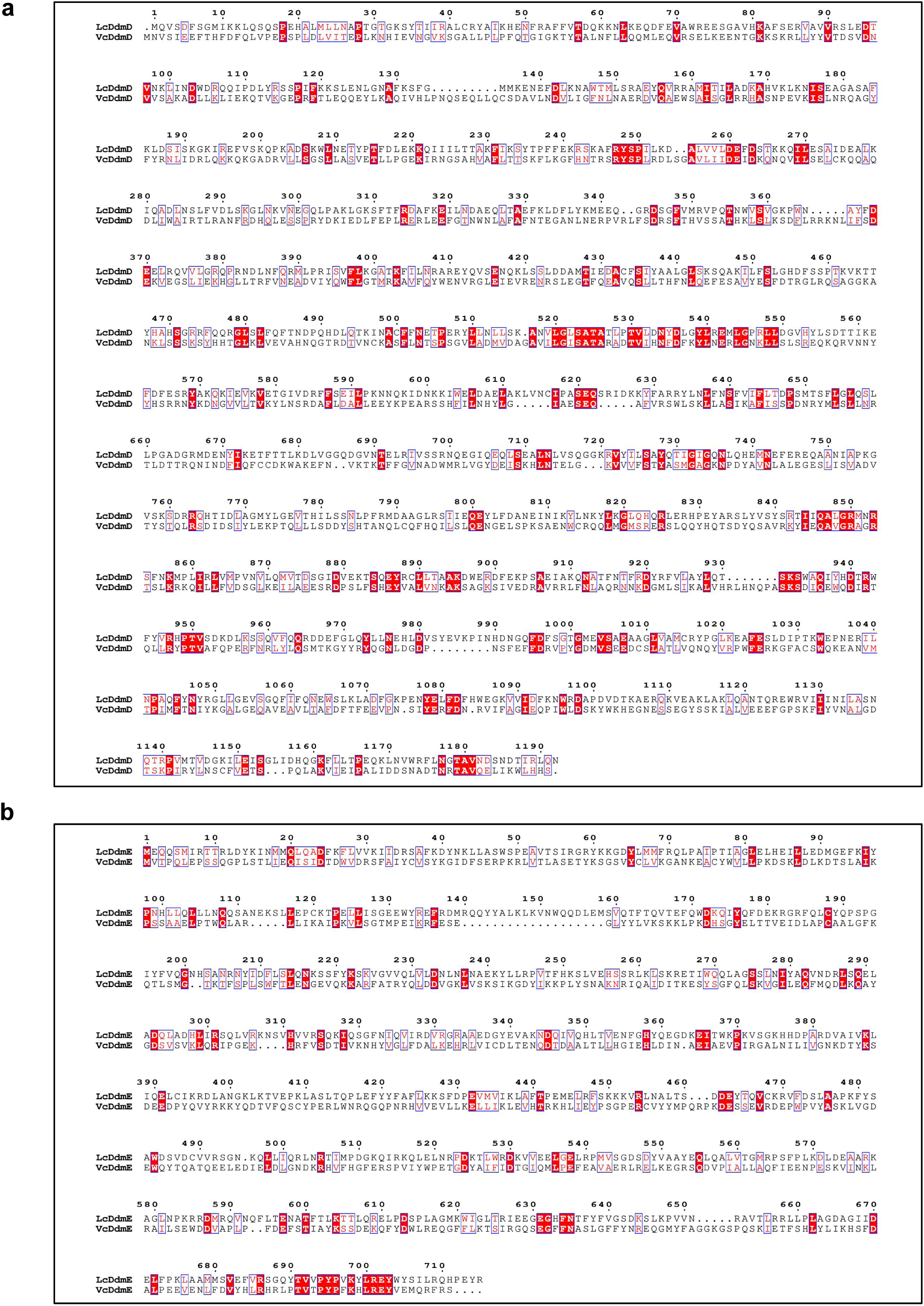
Sequence alignment of DdmD (a) and DdmE (b) from *Lactobacillus casei* and *Vibrio cholerae*.

**Supplementary information, Fig. S10.**
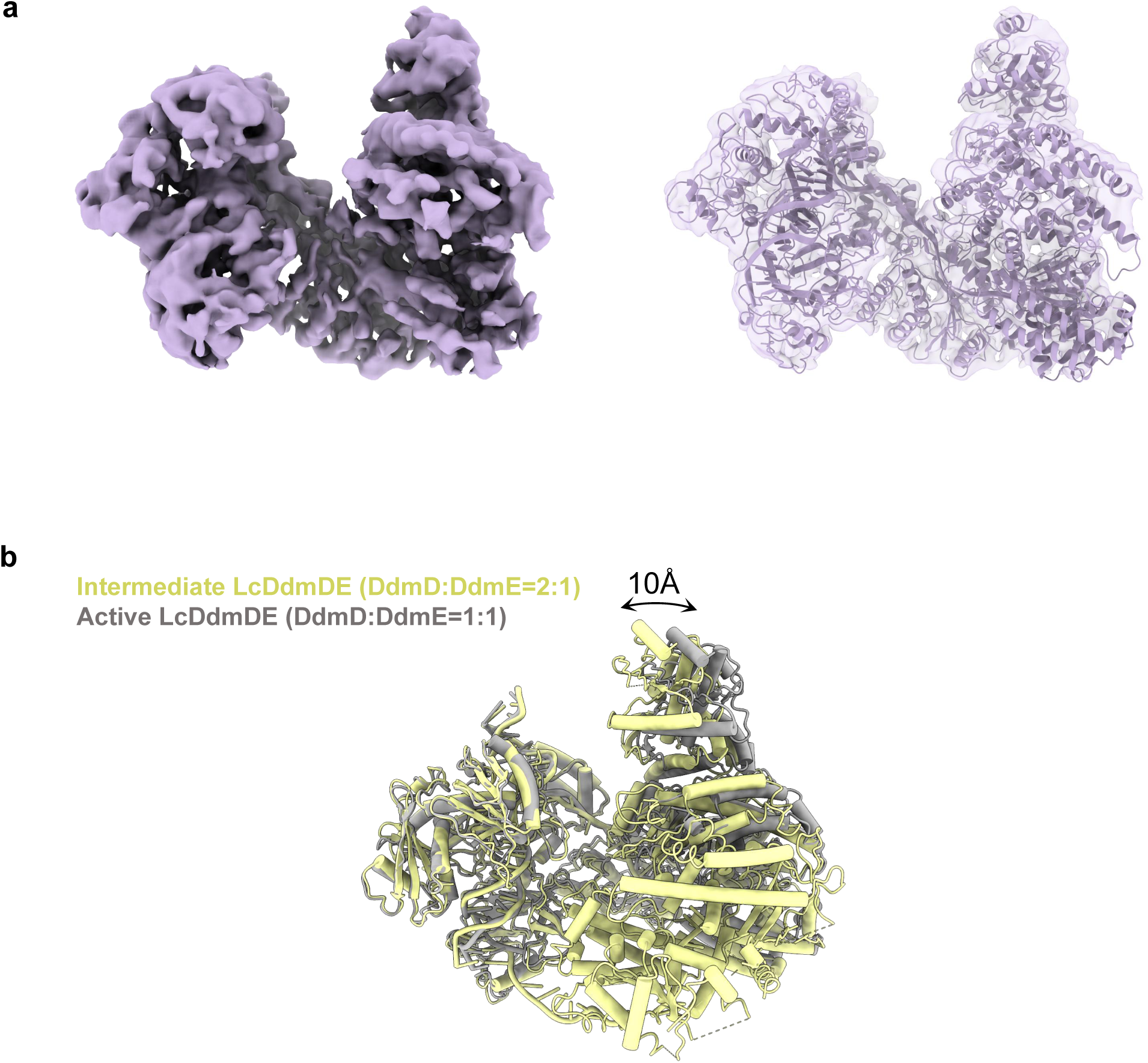
The structure of active LcDdmDE. a. The map and fitted model of DdmDE in active state. b. Structural superimposition of active LcDdmDE (DdmD:DdmE=1:1) onto intermediate LcDdmDE (DdmD:DdmE=2:1).

**Supplementary information, Fig. S11.**
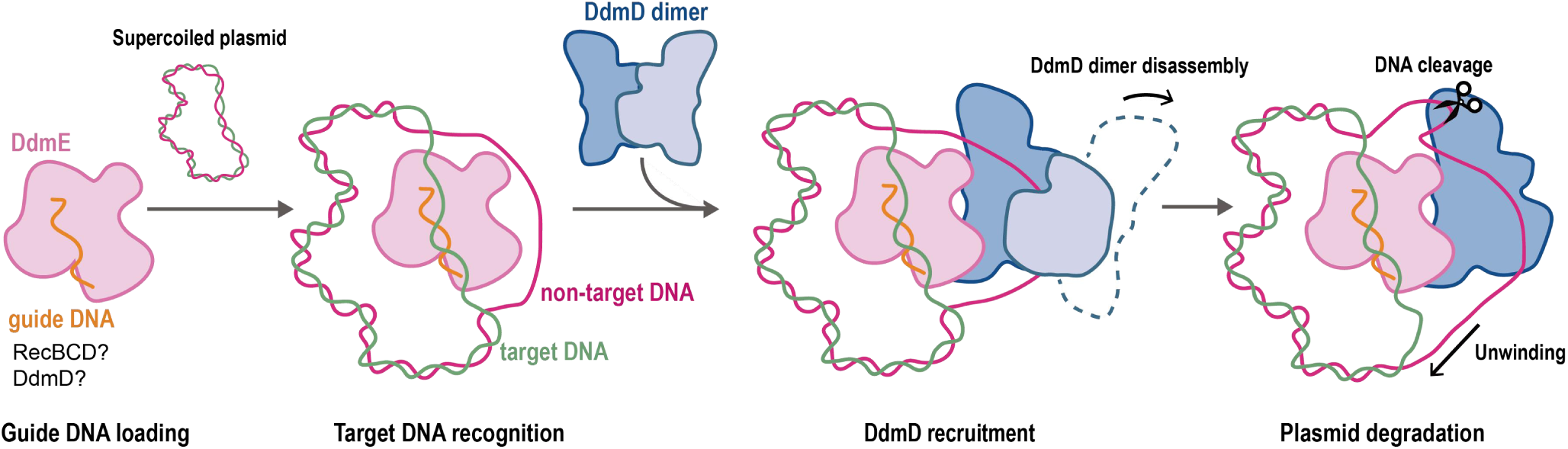
The proposal on mechanism of plasmid degradation by DdmDE. DdmE senses the invasive plasmid directed by guide DNA, which may be generated by RecBCD. DdmD with intrinsic nicking activity may also participate in the first round of guide generation with the assistance of RecBCD. Subsequently, DdmE recruits DdmD dimer onto the bubble region of plasmid via DNA-guided DNA targeting, which facilitates step-by-step dissociation of DdmD dimer in a ssDNA-dependent manner. Ultimately, the DdmD- DdmE heterodimer is activated to unwind and degrade plasmid through the cooperation between helicase and nuclease domains of DdmD.

**Supplementary information, Table S1.**
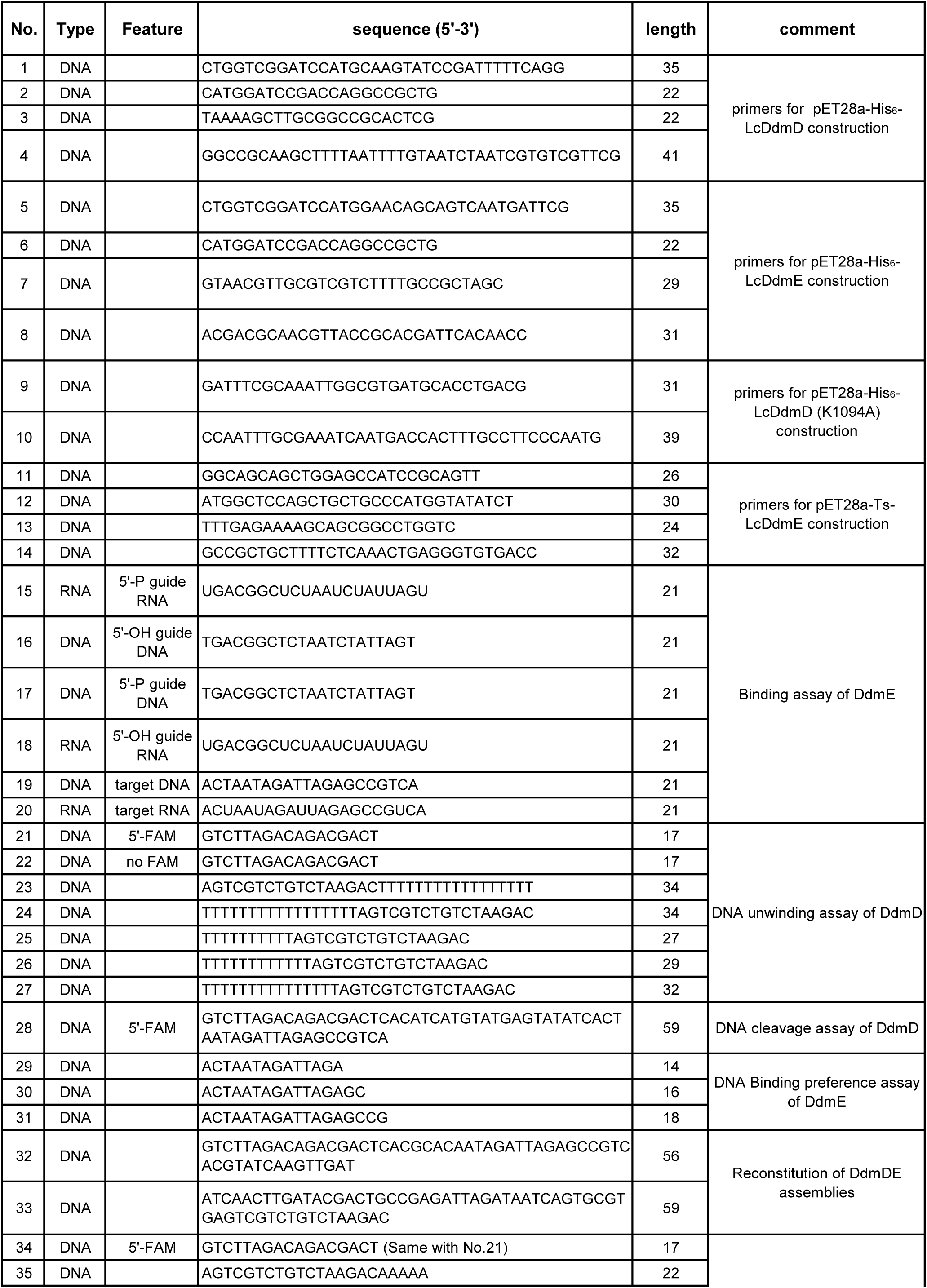
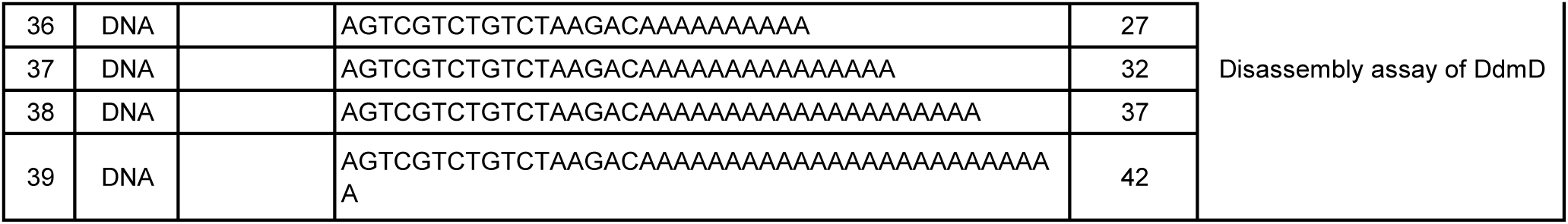
The oligonucleotides used in this study.

**Supplementary information, Table S2.**
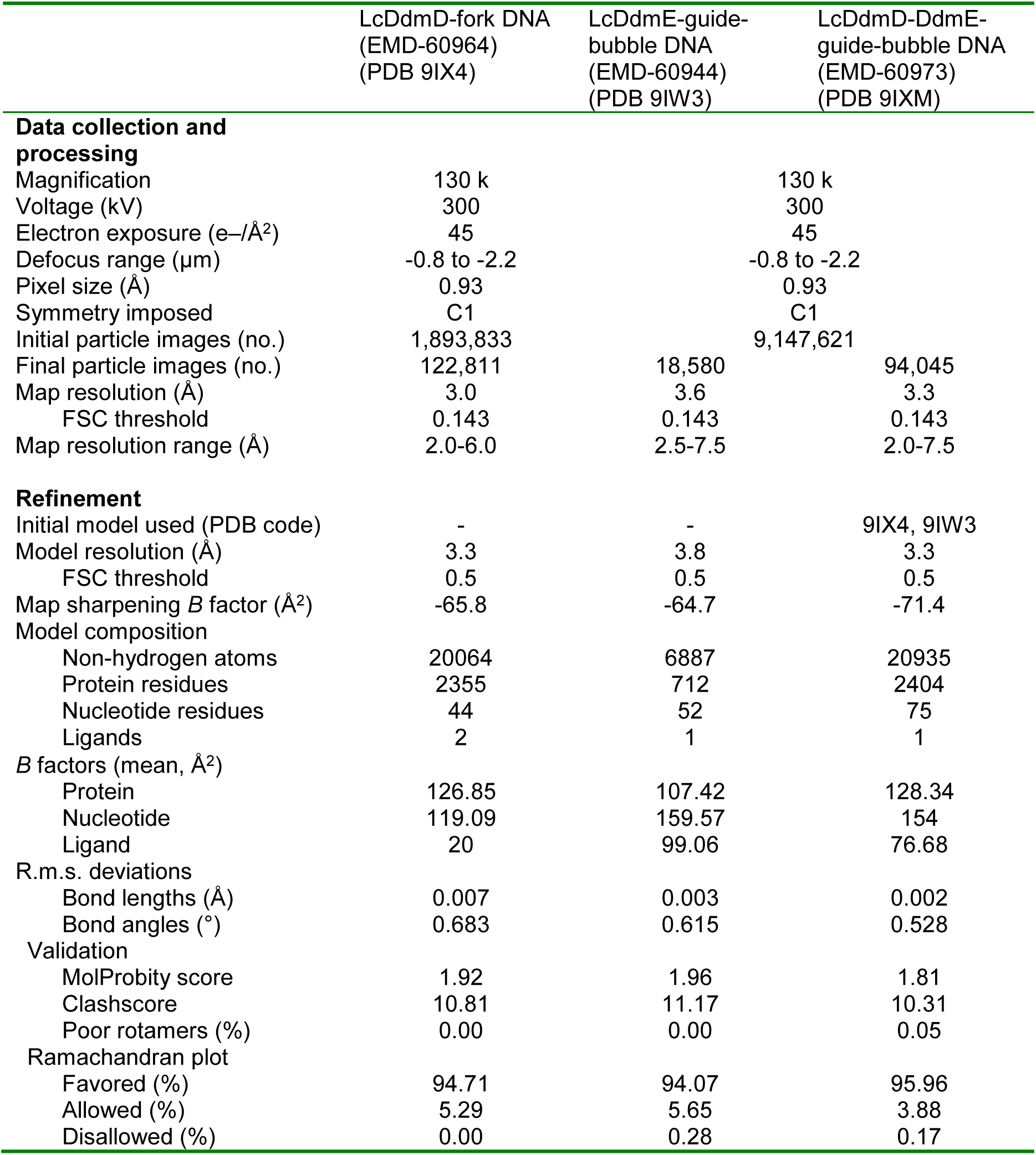
Cryo-EM data collection, refinement and validation statistics.

